# *In silico* classification and identification co-purified protein complexes yield new structures and multiple MSP assembly states

**DOI:** 10.1101/2025.02.04.636456

**Authors:** Qingyang Zhang, Abhinandan Venkatesha Murthy, Carsten Mim

## Abstract

Native protein complexes have garnered interest as targets for structural dissemination. Cryogenic electron microscopy (cryoEM) with its ability to image protein mixtures is the most promising tool to enable structural proteomics. Additionally, image processing has evolved to deal with conformational and compositional heterogeneity. Integrative approaches, namely mass spectrometry in conjunction with cryoEM have made it possible to characterize and identify complex mixtures. However, this comes at a cost of generating models and interpreting mass spectra. Here we present a modified approach that only requires electron micrographs and a computer for unsupervised model building and protein identification. We were able to identify co-purified membrane proteins, resulting in a novel structure and unexpected nanodisc assemblies, which imply direct interaction between membrane proteins and membrane scaffolding proteins.

## Introduction

Visualization of native protein complexes is essential to identify the physiological composition and stoichiometry of biologically relevant protein complexes. In some cases, native complexes show that structures and conformations solved with heterologous expressed proteins are not representative [1]. Protein concentration and heterogeneity of native samples too often restricts structure determination by X-ray crystallography or NMR. However, Cryogenic electron microscopy (cryoEM) has allowed for reconstruction of native complexes early on [2]. But only the advent of single particle analysis (SPA) started a new era in cryoEM and made it routine to solve structures with atomic resolution [3]. Over the last 5 years, the deposition of structures solved by SPA cryoEM has increased almost exponentially (https://www.ebi.ac.uk/emdb/statistics/emdb_entries_year). It has been shown that cryoEM can be a fast structural method, by generating the high resolution structure of a plant ribosome, from the leaf to the final map, in one day [4]. Ever since, structural biology of native samples has become an expanding field [1, 5-7]. Over the last few years, maximum likelihood and deep learning methods have been used successfully to deal with sample heterogeneity, in particular compositional heterogeneity and conformational heterogeneity (e.g. Relion, CryoDrgn, CryoSparc [8-10]), this positions cryoEM as an ideal tool for structural proteomics. One early approach is based on a shotgun proteomics, by fractioning cell lysates through size chromatography and subsequent cryoEM/negative stain EM characterisation of 2D classes and 3D maps [11, 12]. This approach only works with proteins of high natural abundance, e.g. housekeeping or structural proteins. Therefore it is necessary to enrich low copy proteins, say by density centrifugation [4, 7] or by using an affinity chromatography resin with a broad specificity [13]. Immobilized metal affinity chromatography (IMAC) has the ability to co-purify a variety of proteins, in *E*.*coli* [14]. This feat has been used to structurally investigate co-purified complexes [15]. However, structural proteomics for membrane proteins represents a unique challenge, because it requires stabilization of membrane proteins, most often with detergents. Lipid nanodiscs, which are a shell of stabilised lipids around a protein, offer an opportunity to preserve the lipid environment around the protein. The most common nanodisc system uses the membrane-scaffold-protein (MSP) technology and is based on the apolipoprotein A (ApoA) protein, yet other systems are gaining popularity (e.g. reviewed in [16]). A recent study combined the database search of cryoEM-based, *de novo* models with mass spectrometry to identify heterogeneous protein populations. This approach has proven to be a powerful tool [13, 17]. However, this workflow may become tedious because the models must be built, and mass spectrometry data collection and interpretation requires expertise. Lately, AI-assisted model building has opened the avenue for unsupervised model building and protein identification [18, 19].

Here, we present a workflow for the identification of cryoEM maps from a heterogeneous protein mixture. We show that particle classification and unsupervised model building may be sufficient to characterize complex samples with fewer resources. We were able to identify and build models for 3 membrane proteins co-purified from IMAC, one of them represents a new structure. These proteins were reconstituted in MSP2N2 derived nanodiscs and exhibited unusual assembly states. Namely, smaller nanodiscs and non-circular shapes. We identified candidates for interaction sites between membrane proteins and the scaffold proteins.

## Results

### Workflow for the preparation of co-purified membrane proteins in nanodiscs

Initially, this study focussed on the production of a mammalian membrane protein in *E. coli*. Xiang *et al*. demonstrated that the application of an osmotic shock in addition to a cold shock increased the expression of a pentameric ion channel [20]. Therefore, we adapted this protocol with the goal to express the human Zinc activated ion channel. After lysis and removal of debris and cytosolic components, we solubilized the membrane with a high salt buffer (0.9M KCl) to minimize unspecific, ionic interactions. The solubilized membrane proteins were enriched with immobilized metal affinity chromatography (IMAC). Because we were interested in lipid-protein interactions, the enriched protein was incorporated into MSP2N2 nanodiscs, as described for other pentameric channels (e.g. [21]). The sample was further processed with size exclusion chromatography (SEC) and fractions were pooled for characterization. SDS-PAGE showed multiple proteins (Fig. S1). IMAC is known to enrich native *E*.*coli* proteins (e.g. [14, 15]).To characterize these proteins we performed mass spectrometry of the concentrated SEC fractions. We were able to identify many *E*.*coli* specific proteins in our sample (Table 2). Yet, without quantification, the large number of hits makes an identification of the most abundant bands challenging. So, we favoured a cryoEM based approach, which had been used in the past by others [17]. In contrast to the latter study, we wanted to build the model unsupervised as a template for the identification of the proteins through an hidden Markov models (HMM) search [19] with the ModelAngelo software package [18] (Fig. 1).

**Figure 1.**
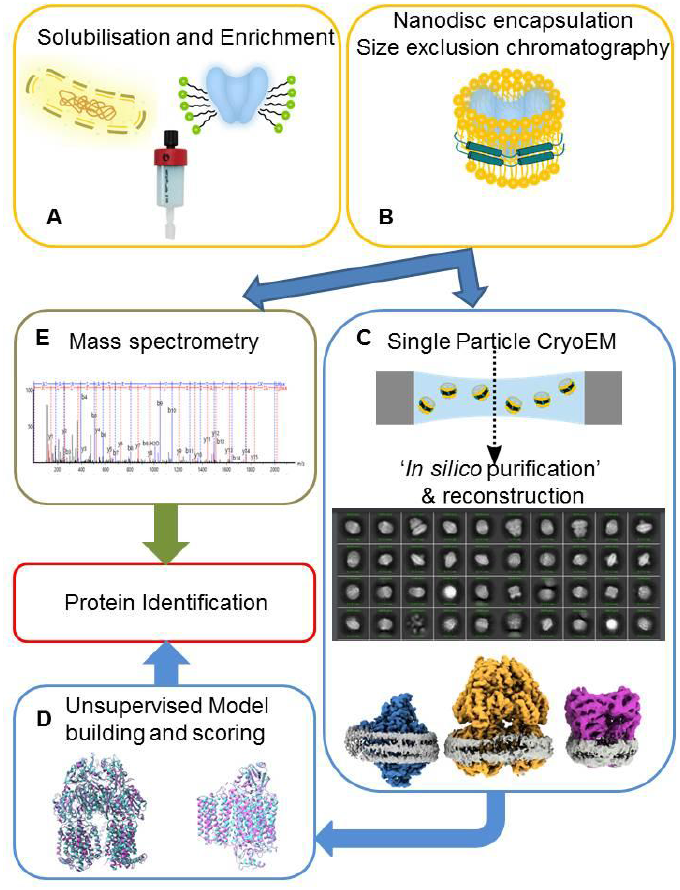
Workflow for the production and *in silico* identification of membrane proteins. (A) *E. coli* membrane proteins are solubilized and enriched with IMAC resin and eluted (B) the enriched proteins are then embedded into nanodiscs with the same scaffolding protein (MSP2N2) and separated by size exclusion chromatography (C) the protein mix is then imaged by single particle cryoEM. The particles are classified and reconstructed separately. (D) The best maps are used to build a model with ModelAngelo [18] without sequence input to generate hits. In addition, a manual search was performed to identify proteins. (E) the same sample that was imaged was analysed via mass spectrometry.

### Unsupervised model building of cryoEM maps can identify proteins

ModelAngelo is a software suite that integrates model building with an HMM search and works most reliably for cryoEM maps with a resolution of 4Å and better [18]. Notably, about 75% of all deposited cryoEM maps in 2024 EM database are 4Å and better (https://www.ebi.ac.uk/emdb/statistics/emdb_resolution_year:retrieved12/2024), which encouraged us to proceed with our approach. We collected almost 18,000 movies. To get an idea of the most prominent proteins in our mixture, we picked particles manually to generate crude templates for template picking. This generated about 3 million particles initially and we classified them into 200 classes (Fig. S2). Among the 200 classes we suspected 3-4 different proteins, which may have been biased by the choice of the template. As a control, we used featureless, circular templates with a diameter of 80 to 180Å (‘blobs’), and we uncovered the same proteins (Fig. S5). We proceeded with separating particles for proteins we deemed similar and applied further rounds of 2D classification and a final *ab initio* reconstruction with multiple classes. This approach yielded in 3 cryoEM maps. To improve particle set we used the best 2D classes of each individual map as a template for picking.

After motion correcting the ‘cleaned up’ particles, we obtained 3 maps with an overall resolution of <4Å, although the local resolution may be lower (Fig. 2). From previous studies [22, 23] and visual inspection, Cytochrome bo(3) ubiquinol oxidase (BO3) was identified as one of the proteins. We chose this as the first map for modelling by ModelAngelo without supplying a sequence and an HMM search. Apart from the non-protein moieties, all subunits were correctly identified. We used the deposited model (pdb ID: 7N9Z,[22]) to model manually and to refine. Even without refinement the backbone trace of unsupervised model (Fig. 2C, magenta) and the manually refined model (Fig. 2C, cyan) are in good agreement. The next we turned to the protein with the largest map. It is an asymmetric trimeric protein and was identified at the multidrug efflux pump subunit ACRB (Fig. 2B). Again, we used a deposited model (pdb ID: 2HRT, [24]) for manual modelling and refinement. The unsupervised model (Fig. 2B, magenta) and the manually refined model (Fig. 2B, cyan) were in good agreement, again. The last map presented a challenge because it represented the smallest protein with a large variation in resolution and areas worse than 4Å in resolution.

**Figure 2.**
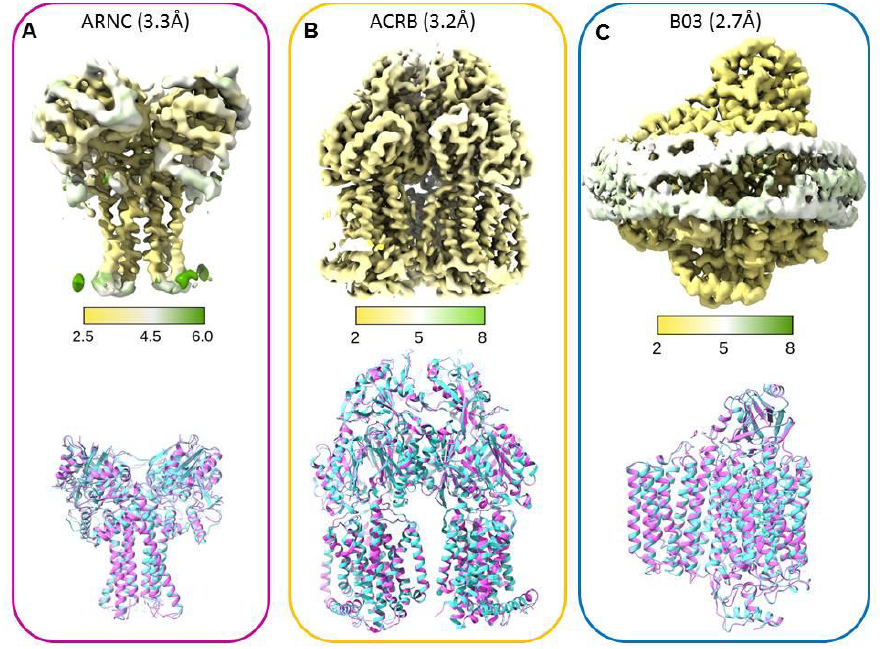
Identification and model building of the reconstructed maps. (A) top: local resolution of the protein density for Undecaprenyl-phosphate 4-deoxy-4-formamido-L-arabinose transferase (ARNC) in Å with an overall resolution of 3.3 Å. bottom: The comparison of the backbone trace from the manually refined model (cyan) with the unrefined model build by ModelAngelo (magenta) shows little deviation. (B) top: local resolution of the map for Multidrug efflux pump subunit ACRB in Å with an overall resolution of 3.3Å. bottom: The comparison of the backbone trace between the manually refined model (cyan) and the ModelAngelo model (magenta). (C) top: local resolution for the Cytochrome bo(3) ubiquinol oxidase subunit 4 map in Å with an overall resolution of 2.7Å. The density for the membrane scaffolding protein is shown as well. The comparison of the backbone trace between the manually refined model (cyan) and the ModelAngelo model (magenta).

Nevertheless, ModelAngelo identified the protein as Undecaprenyl-phosphate 4-deoxy-4-formamido-L-arabinose transferase (ARNC). Because this is a new structure, we used the AlphaFold model (ID: P77757) for refinement. Generally, the unsupervised model (Fig. 2A, magenta) and the manually refined model (Fig. 2A, cyan) agreed well. However, the quality of the model was problematic, because some of the assigned sidechains (e.g. Fig. S6A) and connectivity between some of the helices were incorrect. We were able to verify the identity of all proteins by mass spectrometry (Table. 2).

All target proteins interact directly with MSP2N2 in smaller than expected Nanodiscs We solved the structures of 3 different membrane proteins that were reconstituted in nanodiscs (Fig. 3). All these proteins were of different size and exhibited different nanodisc shapes. This was a surprising finding because we used MSP2N2 as the scaffolding protein for the nanodisc. With the CHARMM-GUI [25] we built a model for a MSP2N2 nanodisc. The theoretical diameter of this nanodisc is 165Å, which was substantially larger than any of the distances in the nanodiscs we observed (ARNC = 65Å, ACRB = 108Å, BO3 = 108Å). The MSP2N2 nanodisc was framed by two helices (Fig. 3 B bottom). In our map we saw variation of this assembly, while the ARNC and ACRB nanodiscs showed two rungs. Some positions of the BO3 nanodisc had 3 rungs of MSP helices, reminiscent to the assembly observed by Roh *et al* [26]. Notably, we saw that the MSP2N2 density is strongest close to the membrane protein (Fig. 3A and B, arrows at zones with lower density). Therefore, we hypothesized that there is an interaction between MSP2N2 and the protein it encapsulates. We searched for residues in our models that are within 5Å of the nanodisc density, which is slightly longer that the contour length of amino acid sidechains [27]. We identified several non-polar and aromatic sidechains in each protein (Fig. 3C-E, Fig. S4). In ARNC, a pair of aromatic residues (W135, F136 Fig 3C) is situated near the nanodisc density. ACRB shows more potential contact sites, through aromatic residues (Y554 and F918, Fig. 3 D; W515, Fig. S6E) and an arginine (R540, Fig. S6E). Last, in BO3 we identified a pair of tryptophan side chains (W34, Fig. 3E, W454 Fig. S6F) and an arginine (R455, Fig. S6F) close to the membrane. We could not generate a separate map for only the MSP2N2 density, good enough to build a model. Yet, all these putative contact sites enforce the idea that these membrane proteins have a more active role in the nanodisc assembly.

**Figure 3.**
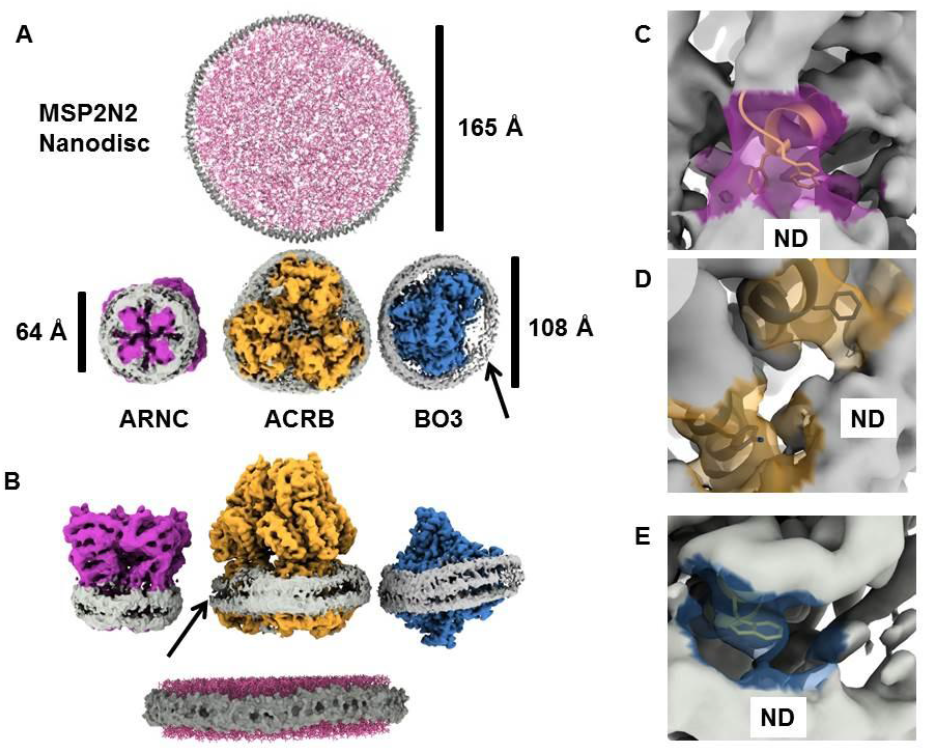
Interactions of membrane proteins with nanodisc scaffolding proteins. (A) top: Top view of an ideal nanodisc formed by MSP2N2 (grey) and lipids (pink) generated by the CHARMM-GUI [25] with a diameter of 165Å and bottom: the nanodisc found in samples. Unsharpened maps of the identified proteins show a different diameter and shape. The ARNC (magenta) nanodisc (grey) has a diameter of around 65Å. The largest extension of the nanodiscs around ACRB (yellow) and BO3 (blue) are similar (108Å) but their shape is different. (B) side view of the structures in A. The scaffolding protein (MSP2N2) in the ideal nanodisc is represented as a space filling model (grey) around lipids (pink). The hydrophobic core is covered by two adjacent MSP2N2 helices (bottom). ARNC and ACRB nanodiscs have densities resemble the two-helix assembly seen in MSP2N2 nanodiscs. B03 exhibits three rungs of helices to form a nanodisc. The scaffold protein density appears to be weakest in areas without the membrane protein in the ACRB and BO3 maps (arrows). (C) putative interaction sites between ARNC (transparent purple) and MSP2N2 (ND). Residues W165 and F136 (yellow) are within 5Å of the MSP2N2 density. (D) ACRB (transparent yellow) contains residues (Y554 and F918) that are within 5Å of the MSP2N2 density (grey). (E) one putative interaction site in BO3 (light blue) showing W34 within 5Å of the MSP2N2 density (grey).

### Structural features of the identified protein complexes

Our purification protocol used high ionic strength to assist solubilisation. We also embedded bacterial proteins into soy lipids. Therefore, we were interested if the structural features of the discovered proteins are altered. In the BO3 map we observed non-protein densities at the position we would expect lipids (Fig. 4A, yellow). Most of the densities were not defined enough to build models for specific lipids. However, we could fit some of the lipids seen in a deposited model (Fig. S6C and D) [22]. We were also able to model the functional groups of the electron transfer pathway, namely the heme moieties and the coordinated metals Zn^2+^ and Cu^2+^ (Fig. 4B). Li *et al* reported that the binding site for the electron carrier ubiquitin (UBQ) is dynamic. Therefore, we performed a 3D variance analysis (3DVA) for the BO3 dataset [10]. The overall structure showed little flexibility, but we observed that the presumed UBQ density varied (Fig. 4C). This could either indicate that some particles lost UBQ or that the position of UBQ is not fixed. In the previous section we showed that all proteins have side chains that are near the nanodisc MSP. There have been concerns that the nanodisc may change the structure of the embedded membrane proteins [28, 29]. ACRB is a pump whose subunits cycle through 3 conformations: ‘loose’ (L) and ‘tight’ (T) are structural similar and have substrate binding sites open to the periplasm; ‘open’ (O) has the substrate binding site closed to the periplasm [30]. We performed a 3DVA with the ACRB particle set. When we separate 4 clusters, we observe that 3 clusters in an asymmetric LTO state (Fig. 4D for the L protomer conformation, and 4E for the O protomer conformation, the T state was omitted for simplicity, movie M1)[24]. The LTO state was modelled for our reconstruction (Fig. 2B). Yet, one cluster represents a symmetric state that we believe is in an all ‘loose’ state, based on the fitting of the ‘loose’ subunit. Last, we solved the structure of ARNC embedded in the membrane, unlike the structures of homologues, which were solved in detergent [31, 32]. The helices closest to the membrane (amino acids: 215-228 and 135-153) are both amphipathic helices (Fig. S4B) and the map suggest a placement of these helices in one leaflet. These helices comprise the membrane adjacent part of the substrate binding pocket in ARNC homologues [31, 32].

**Figure 4.**
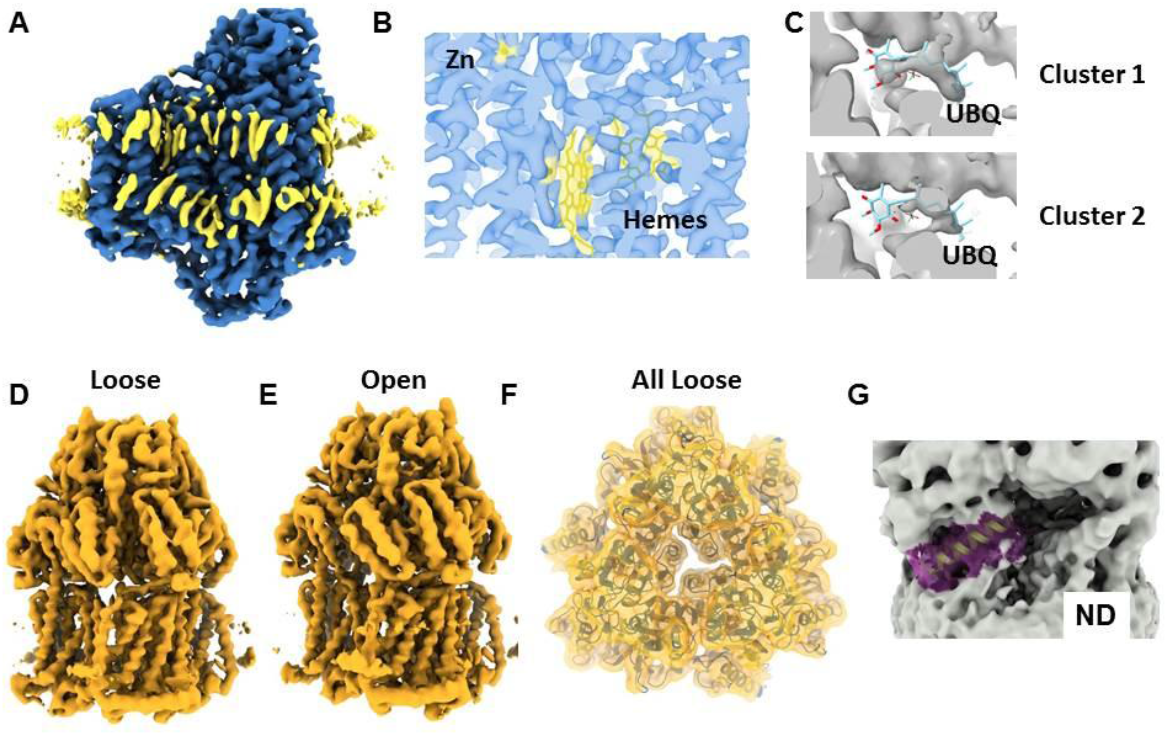
Structure-function features of the characterized proteins. (A) Non-protein densities (yellow) in the BO3 map at the position of the bilayer’s hydrophobic core. (B) Density for the protein (transparent blue) and non-protein moieties (transparent yellow) in BO3. (C) 3DVA analysis of the density (grey) for the electron carrier ubiquitin (UBQ, stick representation) in BO3. Three clusters were investigated, and the UBQ-associated density varies. (D) and (E) 3DVA analysis of the ACRB dataset (See movie M5) shows several conformations of the efflux pump, ‘Loose’, ‘Open’. The ‘Tight’ conformation is not shown here for simplicity but modelled. (F) one of the clusters from the 3DVA shows an all ‘Loose’ conformation (G) Putative substrate binding domain of ARNC (yellow in purple envelope) and its proximity to the membrane.

## Discussion

Structural biology has long relied on pure samples with engineered proteins that enhance stability to either limit aggregation at high concentrations or to endure the long timescales necessary to grow crystals. With high resolution cryoEM limitations on the sample became less restrictive, because cryoEM is an imaging technique. As a result, solving the structures native proteins and complexes has become more common (e.g. [1, 4, 12, 33]). However, some of studies still require the use of affinity proteins like nanobodies. The ‘build and retrieve’ approach worked with an IMAC resin, yet supervised (*de novo*) model building or *a priori* knowledge of the reconstructed map is part of that workflow [17]. Here we present and alternative for an unbiased workflow to identify reconstructed maps of unknown proteins. Our workflow suggests that the mass spectrometry is not strictly necessary, which may make our approach more accessible because it reduces the cost and time for mass spectrometry data collection and interpretation. We have not tested the workflow with a map with less than 4Å in resolution, but the ARNC map has many areas where the resolution is less than 4Å and the assignment of residues in some areas is error prone. Yet, there were enough modelled fragments that allowed a correct identification. We imagine that the identification of protein isoforms may prove difficult at lower resolutions and mass spectrometry may help in those cases. Notably, the combination of proteins we identified has been unique. BO3 seems a common protein co-purified with IMAC [17, 22, 23]. However, ACRB and ARNC are novel proteins appearing as co-purified proteins. While ARNC represents only a small fraction of the particles, ACRB is a prominent component. This may be due to the chosen templates but even ‘blob’ based picking resulted in a population of ACRB/ARNC in the 2D classification (Fig. S5). It appears that each ‘non-specific enrichment’ purification may result in different protein populations and may depend on the ionic strength, used detergent, expression protocol and other factors. Interestingly, ACRB and ARNC are proteins that are overexpressed in response to antibiotics.

Our study also provided insight into interactions in and assembly of MSP based nanodiscs. We demonstrate MSP2N2 can accommodate different nanodisc sizes and shapes. The size of the ARNC map implies that one MSP2N2 molecule is enough to form a nanodisc. This deviates from the common assumption that 2 MSP molecules are necessary [34]. Although it has been reported that 3 MSP molecules can form a nanodisc [35], the smaller size of the BO3 nanodisc makes it unlikely, despite the observation of 3 rungs of helices. The flexibility in nanodisc formation has only been observed with SaliPro nanodiscs [36]. Like SaliPro nanodiscs, we saw that the MSP2N2 density is stronger around the membrane protein. This suggests direct interactions between the embedded protein and the scaffolding protein. We suggest that the interaction with the membrane protein enables diverse assemblies and may steer the lipid-to-protein ratios in nanodisc, to the point that no detergent or lipids are necessary for the nanodisc formation [37]. These findings are somewhat in conflict with standardized protocols for MSP-based nanodisc formation[34]. The lack of lipids may be detrimental to the function of the protein, because the lateral pressure in small nanodiscs is higher [29] resulting in conformational bias depending on the nanodisc [28]. Because we observed no density in any binding sites, we presume it is an apo state of ACRB. Yet, we observe an asymmetric LTO configuration, where at least one protomer (T) should be stabilized by substrate. Notably, the asymmetric LTO state has been observed in narrow MSP nanodiscs with or without substrate, and the binding mode of the substrate in nanodiscs is different from the one seen in a crystal [38]. However, we could identify a population in a presumed resting state in (all L) [39]. At this point it is unclear if the nanodisc arrests ACRB in a state where the subunits cycle through a transport cycle or if the nanodisc stabilizes the resting state.

Taken together, this study has shown that our approach is feasible and leads to novel insights. Yet, more studies with more complex protein mixtures, and a wider range of cryoEM map resolutions may be necessary to determine how robust our approach is.

## Materials and Methods

### Protein expression

BL21-Tuner cells (Novagen) were transformed with XXXX in pET22b (pET22b was a gift from EMBL protein production core). For expression, cells were grown in TB medium at 37°C until OD_600_ of 0.72. Then the culture was pelleted at 6000g, and the medium was exchanged to TB media supplemented with 250mM sorbitol and equilibrated at 18° C for 45mins. At an OD_600_ of 0.75 expression was induced with 0.5mM IPTG at 20° C overnight.

### Solubilisation and Enrichment

Cells were resuspended (50mM Tris, pH 7.2 at RT, 500mM NaCl, 5% glycerol) and cOmplete protease inhibitor cocktail tablets (Roche) were added. For lysis 1uL per mL of 100mg/ml lysozyme (Sigma) was added and incubated at 4C for 1h. Cells were lysed by sonication (Thermo Fisher Scientific) 50% amplitude, 10mins, 5s ON ; 15s OFF. Debris was removed by centrifugation at 10000 g for 30 mins. Membranes were harvested at 108000g for 1h, 4C. The membranes were homogenised in 50mM HEPES, 900mM KCl, 5% glycerol and one protease inhibitor cocktail tablet. Membranes were solubilized with 1% n-Dodecyl β-D-maltoside (DDM, Anatrace)/0.2% Cholesterol Hemi Succinate (CHS, Sigma) 3-4h at 23°C. Non-solubilized material was removed at 108000g for 30mins at 4°C. The supernatant was incubated with NiNTA resin overnight at 4°C. The resin was washed with homogenisation buffer with w/v 1%DDM/0.2%CHS containing 10mM imidazole. The protein was eluted in fractions with the Wash buffer and 250-300mM imidazole.

### MSP encapsulation and size exclusion

The elution fractions were pooled, and the buffer was exchanged to 50mM HEPES (pH=7.5), 300mM KCl w/v 1%DDM/0.2%CHS with PD10 columns (Cytiva) according to manufacturer’s recommendations. After protein concentration determination (OD_280_) 1:65 (protein to lipid molar concentration) of Soy Polar lipids (Avanti) were added and incubated for 30 min. Then 1:5 (pentameric protein to MSP molar concentration) of MSP2N2 was added and incubate for 30-40mins. 50mg of wet/activated S2 biobeads (BioRad) were added and incubated for 3-4h. If detergent was not completely removed, another 50mg of biobeads were added and incubated 3h or overnight. (add more biobeads as required). After detergent removal biobeads were removed and the remaining solution was incubated with NiNTA (G-Biosciences) for 1.5-3h. The beads were washed (50mM HEPES (pH=7.5), 300mM KCl and 10mM imidazole) and the proteins eluted (50mM HEPES (pH=7.5), 300mM KCl and 250mM imidazole). The pooled fractions were concentrated subjected to SEC (ÄktaExplorer, Cytiva) with a Superose 6 column (Cytiva) with the SEC buffer: 50mM HEPES (pH=7.5), 150mM NaCl and 5mM EGTA.

### CryoEM sample preparation and cryoEM data collection

SEC Fractions were pooled and concentrated to 4.6mg/ml. Quantifoil 2/2 Cu 300 grids were glow discharged for 60 s at 20mA and 3μl of the sample was applied to the grids. The grids were blotted for 7 s, at 6° C, 100% humidity with a Vitrobot Mark IV (Thermo Fisher Scientific). The sample was imaged on a Krios (Thermo Fisher Scientific) equipped with a K3 camera in super resolution mode (Gatan). The movies were collected at a magnification of 105k a total dose of 50 e^-^/Å^2^ over 40 frames. The nominal magnification was 130,000X. The total dose was 50 e^−^/Å^2^. The defocus was varied between –1 and –2.8 μm. Data were collected using the EPU software (Thermo Fisher Scientific).

### Cryo-EM single particle analysis

All data were processed using CryoSparc version 4.1-4.5 (Structura Biotechnology Inc., Toronto, ON, Canada) [40]. The raw movies were motion-corrected using patch motion correction software. The contrast transfer function (CTF) was estimated using the software’s patch CTF estimation. The micrographs were curated based on the CTF fit resolution, motion distance, and ice quality. The particles were manually selected from a small subset of images and classified in 2D. 2D classes with a discernible shape were used as coarse templates for picking. The particles were down sampled and subjected to multiple rounds of 2D classification. 2D classes that shared similarity were selected and further classified. The best 2D classes were used for *Ab initio* map generation. The obtained maps were refined with non-uniform refinement. To generate a more specific particle set we used 2D classes of particles sets for each reconstructed protein as templates. Due to the small total number of particles for ARNC, we used the best particles set as a training set for Topaz [41] to pick particles. All different protein particles were curated using two-dimensional classification.

The remaining particles were further sorted using *Ab Initio* map generation. After refinement, particles were motion corrected before the final non-uniform refinement [42]. For ACRB, we used one of the asymmetric volumes from the 3DVA as an input volume for the refinement, due to the low-resolution intrinsic symmetry of the ACRB trimer, resulting poorly resolved protomers, even with C1 symmetry applied. For ARNC, local refinement with a mask including only the ARNC density was performed. To control for template picking bias, circular, featureless templates with 80-180Å diameter were used for picking and the 2D classes were analysed for ACRB, BO3 and ARNC particles.

### 3DVA and refinement

For CryoSparc 3DVA [10], the refined particles expanded symmetrically (C3) for ACRB and not expanded for B03. The focusing mask was based on a model we created. For visualisation the volumes we generated in ‘cluster mode’.

### Model building, validation, and structure analysis

For ACRB pdb ID: 2HRT, for BO03 pdb ID: 7N9Z (with lipid densities removed) and for ARNC the AlphaFold prediction of *E. coli* ARNC (P77757) was downloaded from AlphaFold-DB [43] and rigidly fitted into the cryo-EM map using ChimeraX version 1.6.1 [44]. Sequences and side chains of the atomic model that were not supported by any density were deleted. The fitted models were refined against the unsharpened cryo-EM map by interactive molecular dynamics flexible fitting (iMDFF) using ISOLDE version 1.6.0 [45] within ChimeraX[44]. A post-processed map generated by deepEMhancer [46] using tightTarget weights was used as a visual aid for interactive model refinement in ISOLDE (but was not used to drive MDFF). We used a strategy outlined in [47]. Each residue was visually inspected at least once. The resulting model for ACRB were submitted for a final real-space refinement using phenix.real_space_refine from the Phenix Suite version 1.20.1-4487 [48] using the settings file generated by ISOLDE. The models for ARNC and BO3 were refined with ServalCat [49]. All model refinement was done against the unsharpened cryoEM maps. All images were generated using the ChimeraX.

### In-gel Protein Digestion and Mass Spectrometry

Protein bands were excised manually from gels and in-gel digested. Gel pieces were destained following the manufacturer’s description. Proteins then were reduced with 0.25 μL of 500 mM dithiothreitol for 45 min at 37°C and alkylated with 0.75 μL of 500 mM iodoacetamide for 30 min at room temperature followed by digestion with 0.5 μg sequencing grade trypsin (Promega) in 50 mM ammonium bicarbonate at 37°C overnight. The tryptic peptides were extracted with 1% formic acid in 2% acetonitrile, followed by 50% acetonitrile twice. The liquid was evaporated to dryness on a vacuum concentrator (Eppendorf).

The reconstituted peptides in solvent A (2% acetonitrile, 0.1% formic acid) were separated on a 50 cm long EASY-spray column (Thermo Fished Scientific) connected to an Ultimate-3000 nano-LC system (Thermo Fisher Scientific) using a 60 min gradient from 4-26% of solvent B (98% acetonitrile, 0.1% formic acid) in 55 min and up to 95% of solvent B in 5 min at a flow rate of 300 nL/min. Mass spectra were acquired on a Q Exactive HF hybrid Orbitrap mass spectrometer (Thermo Fisher Scientific) in m/z 375 to 1500 at resolution of R=120,000 (at m/z 200) for full mass, followed by data-dependent HCD fragmentations from 17 most intense precursor ions with a charge state 2+ to 7+. The tandem mass spectra were acquired with a resolution of R=30,000, targeting 2×105 ions, setting isolation width to 1.4 Th and normalized collision energy to 28%.

Acquired raw data files were analyzed using the Mascot Server v.2.5.1 (Matrix Science Ltd., UK) and searched against SwissProt protein database with E. coli species selection.

Maximum of two missed cleavage sites were allowed for trypsin, while setting the precursor and the fragment ion mass tolerance to 10 ppm and 0.02 Da, respectively. Dynamic modifications of oxidation on methionine, deamidation of asparagine and glutamine and acetylation of N-termini were set. Initial search results were filtered with 5% FDR using Percolator to recalculate Mascot scores. Protein identifications were accepted if they could be established at greater than 95.0% probability and contained at least 2 identified peptides.

## Acknowledgments

Protein identification was carried out by the Proteomics Biomedicum core facility, Karolinska Institutet (https://ki.se/en/research/proteomics-biomedicum-core-facility). Data were collected at the Cryo-EM Swedish National Facility funded by Knut and Alice Wallenberg, Family Erling Persson, Kempe Foundations, SciLifeLab, Stockholm University, and UmeÅ University.

## Funding

This study was supported by the Carl Trygger Foundation grant (CM).

## Data availability

CryoEM maps were deposited at the Electron Microscopy Data Bank with the accession codes EMD-XXXXX (ACRB), EMD-XXXXX (BO3) and EMD-XXXXX (ARNC). The models were deposited at the Protein Data Bank with the accession codes XXXX (ACRB), XXXX (BO3) and XXXX (ARNC).

## Author contribution

Purification and sample preparation: AVM, QZ

Data processing and model building: CM, QZ, AVM

Data interpretation: CM, QZ, AVM

Manuscript preparation and revision: CM, QZ, AVM

Competing interests: Authors declare that they have no competing interests..

---

**Table 1.** Image processing and model building statistics.

|  | ACRB | BO3 | ARNC |
| --- | --- | --- | --- |
| Magnification | 130000 | 130000 | 130000 |
| Voltage(kV) | 300 | 300 | 300 |
| Electron exposure (e-/Å2) | 50 | 50 | 50 |
| Defocus range (-µm) | 1-2.5 | 1-2.5 | 1-2.5 |
| Pixel size (Å) | 0.66 | 0.66 | 0.66 |
| Symmetry imposed | C1 | C1 | C4 |
| Initial particle images (no.) | 396,209 | 794,072 | 544,237 |
| Final particle images(no.) | 82,720 | 160,179 | 48,142 |
| Map resolution (Å) | 3.27 | 2.72 | 3.27 |
| FSC curve | normal | auto-tightening | normal |
| FSC threshold | 0.143 | 0.143 | 0.143 |
| Map resolution range (Å) | 1.4-40.4 | 1.4-44.4 | 2.9-34.5 |
| Initial model used (PDB code) | 2HRT | 7N9Z | P77757 (AlphaFold) |
| Map sharpening B factor (Å2) | refinement against unsharpened map |  |  |
| Model composition |  |  |  |
| Non-hydrogen atoms | 23576 | 15043 | 9540 |
| Protein residues | 3104 | 1198 | 1204 |
| Ligands | 0 | 4 | 0 |
| subunits | 3 | 4 | 4 |
| R.m.s. deviations |  |  |  |
| Bond lengths (Å) | 0.01 | 0.009 | 0.011 |
| Bond angles (°) | 0.864 | 1.993 | 1.771 |
| Validation |  |  |  |
| Clashscore | 1.43 | 0.58 | 11.17 |
| Poor rotamers (%) | 0.16 | 0.20 | 0.38 |
| Ramachandran plot |  |  |  |
| Favored (%) | 98.22 | 98.16 | 98.64 |
| Allowed (%) | 1.78 | 1.84 | 1.36 |
| Disallowed (%) | 0.00 | 0.06 | 0.00 |

## Notes

### Competing Interest Statement

The authors have declared no competing interest.

